# Exploratory spatial peptidomic profiling during incubation of drug seeking following cocaine plus alcohol self-administration in young adult rats

**DOI:** 10.64898/2026.08.03.742404

**Authors:** Natalia Puig, Carlos Alberto Castillo-Sarmiento, Lucía Garrido-Matilla, Alberto Marcos, Juan Ramón Peinado, Yoana Rabanal-Ruiz, Daniel Saiz-Sánchez, Enrica Spano, Carlos Vera Fernández, Inmaculada Ballesteros-Yáñez, Emilio Ambrosio

## Abstract

**Background:** Concurrent cocaine and alcohol use is one of the most prevalent forms of polysubstance consumption and is associated with poorer clinical outcomes than cocaine use alone. However, the regional molecular adaptations induced by combined exposure remain poorly understood. Here, we used matrix-assisted laser desorption/ionization imaging mass spectrometry (MALDI-IMS) to characterize peptide/protein alterations in addiction-related brain regions following cocaine and cocaine–alcohol self-administration.

**Methods:** Young adult male and female Wistar rats underwent intravenous self-administration of saline, cocaine (1 mg/kg/infusion) or cocaine plus ethanol (1 mg/kg cocaine and 133 mg/kg ethanol per infusion), followed by extinction of drug-seeking behaviour. Coronal brain sections containing the anterior cingulate cortex (ACC) and ventral hippocampus (vHPC) were analysed by MALDI-IMS. Differential molecular features were identified using an exploratory statistical approach (FDR q < 0.20) and subsequently subjected to MS/MS analysis.

**Results:** The ACC exhibited a substantially greater number of treatment-associated molecular alterations than the vHPC, suggesting a higher regional susceptibility to cocaine-induced molecular remodelling. Several molecular features were shared between the cocaine and cocaine–alcohol groups, indicating persistent cocaine-driven neuroadaptations. In contrast, additional signals were selectively associated with combined cocaine–alcohol exposure, while others present after cocaine alone were absent following alcohol co-exposure, supporting a modulatory effect of alcohol on specific cocaine-induced molecular responses. Overall, combined exposure generated a distinct regional molecular profile rather than simply reproducing the effects of cocaine alone.

**Conclusions:** This exploratory study demonstrates that MALDI-IMS enables the identification of region-specific peptide/protein alterations associated with cocaine and cocaine–alcohol exposure while preserving their spatial distribution within the brain. These findings highlight the ACC as a particularly responsive region and provide a framework for future studies aimed at validating molecular pathways involved in cocaine–alcohol polysubstance use.

## Introduction

Drug addiction is a chronic neuropsychiatric disorder characterized by persistent neurobiological adaptations that disrupt reward processing, executive control, emotional regulation and memory-related circuitry (Koob and Volkow, 2016). Chronic cocaine exposure induces widespread molecular and cellular remodelling within corticolimbic brain regions, including alterations in dopaminergic neurotransmission, synaptic plasticity, neuroinflammatory signalling and epigenetic regulation (Stewart et al., 2021). These long-lasting neuroadaptations are thought to underlie craving, compulsive drug-seeking behaviour and the high vulnerability to relapse that characterizes cocaine use disorder (Koob and Volkow, 2016).

Within the corticolimbic circuitry involved in addiction, the anterior cingulate cortex (ACC) contributes to executive control, decision-making, emotional regulation and the processing of drug-associated cues. Structural, functional and molecular alterations in this region have been described following repeated exposure to psychostimulants and alcohol and have been associated with impaired inhibitory control and cue-induced craving (Goldstein and Volkow, 2011; Paludetto et al., 2024). The ventral hippocampus (vHPC), through its functional connections with the prefrontal cortex, amygdala and nucleus accumbens, participates in contextual and emotional memory processes that influence drug-seeking behaviour. Neuroadaptations in the vHPC may therefore contribute to the persistence and retrieval of drug-associated contextual memories and to vulnerability to relapse (Barr et al., 2020; Rogers and See, 2007).

Polysubstance use represents a major challenge in addiction research, as cocaine is frequently consumed in combination with alcohol (Pergolizzi et al., 2022). Their simultaneous use results in the hepatic formation of cocaethylene, an active metabolite with a longer half-life than cocaine that may contribute to the enhanced toxicity and reinforcing effects associated with this pattern of consumption (Jatlow, 1993). Beyond this pharmacokinetic interaction, cocaine–alcohol co-exposure may induce neurobiological adaptations that differ from those produced by cocaine alone, involving changes in neurotransmission, oxidative stress, cellular metabolism and molecular regulation (Pennings et al., 2002; Say et al., 2022). Nevertheless, the molecular consequences of their combined consumption within corticolimbic brain regions remain incompletely characterized.

Our collaborative research has progressively characterized the neurobiological consequences of combined cocaine and alcohol exposure using increasingly translational experimental models. Initial studies based on passive drug administration demonstrated that the combination of both substances induces specific molecular alterations that differ from those produced by cocaine or alcohol alone, affecting neurotransmitter systems, including GABAergic, glutamatergic and endocannabinoid pathways (Marcos et al., 2022). These findings were subsequently extended by spatial peptidomic analyses using MALDI imaging mass spectrometry, which revealed region-specific peptide/protein signatures across several corticolimbic structures following combined cocaine and alcohol exposure (Marcos et al., 2025). Building upon these findings, we subsequently implemented intravenous self-administration paradigms, a model that more closely reproduces the voluntary patterns of drug intake observed in humans. Using transcriptomic and metabolomic approaches, these studies consistently showed that cocaine–alcohol co-use generates a distinct neurobiological signature associated with alterations in neurotransmission, cellular metabolism and molecular plasticity, identifying persistent and sex-dependent alterations in brain metabolite profiles and in the expression of genes related to the endocannabinoid, glutamatergic and GABAergic systems (Lucia Garrido-Matilla et al., 2026; Lucía Garrido-Matilla et al., 2026; Marcos et al., 2023).

Repeated drug exposure induces progressive and potentially long-lasting neuroadaptations that promote the transition from voluntary drug use to compulsive consumption and contribute to the persistent risk of relapse (Koob and Volkow, 2016). These adaptations affect a distributed network of interconnected brain regions involved in reward processing, reinforcement learning, executive and inhibitory control, emotional regulation and drug-associated memory. The ventral and dorsal striatum participate, respectively, in reward-driven behaviour and in the progressive development of habitual and compulsive drug seeking (Clarke and Adermark, 2015). In parallel, dysfunction of the prefrontal cortex, including the anterior cingulate cortex, impairs decision-making and inhibitory control and may facilitate continued drug intake despite its adverse consequences (Goldstein and Volkow, 2011). The amygdala and hippocampus contribute to the emotional and contextual components of drug-associated memories, which can persist during abstinence and promote cue- and context-induced drug seeking (Koob, 2009; Kutlu and Gould, 2016). Studying molecular alterations across these corticolimbic and striatal regions is therefore essential to understand how repeated drug exposure produces enduring loss of control over consumption and vulnerability to relapse.

MALDI imaging mass spectrometry provides a valuable approach for investigating drug-induced neuroadaptations by enabling the label-free detection of a broad range of molecular species while preserving their anatomical distribution within brain tissue. This capacity allows region-specific peptide, protein, lipid and metabolite alterations to be examined without the need for predefined molecular targets, thereby complementing conventional biochemical and omics approaches (Aichler and Walch, 2015; Shariatgorji et al., 2014). Although MALDI-IMS has been widely used to detect and map drugs and their metabolites in biological tissues, particularly in pharmacological, toxicological and forensic applications (Granborg et al., 2022), its use to characterize drug-induced alterations in endogenous peptide and protein profiles remains comparatively uncommon. Studies applying this approach to polysubstance exposure are even scarcer. Our previous work provided initial evidence that combined cocaine and alcohol administration produces distinct spatial peptide/protein profiles across corticolimbic brain regions (Marcos et al., 2025). The present study extends this contribution by applying MALDI-IMS to a cocaine–alcohol self-administration model and to the subsequent incubation of drug seeking.

Despite the growing recognition that cocaine and alcohol are among the most frequently co-consumed drugs, the molecular mechanisms underlying their combined effects on the brain remain poorly understood. This limitation is even more evident for spatial molecular approaches such as MALDI imaging mass spectrometry, which have only exceptionally been applied to characterize the endogenous peptide and protein alterations associated with combined drug exposure. Building upon our previous demonstration that passive cocaine–alcohol administration induces region-specific peptide/protein alterations, the present study extends this approach to an self-administration model, providing a more translational framework to investigate the spatial molecular adaptations associated with cocaine–alcohol intake and the subsequent incubation of drug seeking. Therefore, the aim of the present study was to characterize the spatial peptide/protein alterations induced by combined cocaine and alcohol self-administration in rats during the incubation of drug seeking using MALDI imaging mass spectrometry and tandem mass spectrometry-based peptide identification.

## Methods

### Animals

Forty-eight adult male (n=24) and female (n=24) Wistar rats (Charles River Laboratories, Lyon, France), 52 days old at the beginning of the treatment, were housed in same-sex groups of 3 to 4 individuals per cage, with a constant temperature (22 ± 2 °C) and a 12 h light/dark cycle. Food and water were available *ad libitum*. Rats were housed and handled in the animal facility at the Faculty of Psychology, National University for Distance Learning (UNED; Madrid, Spain) in accordance with European Union legislation on the Protection of Animals Used for Scientific Purposes (Directive 2010/63/EU) and with the approval of the Bioethics Committee of the UNED.

### Experimental design

The present study employed an independent cohort of Wistar rats following the same intravenous self-administration protocol previously developed and validated (Marcos et al., 2023). Treatments were randomly assigned according to the experimental design shown in **Fig. 1**. The study included three independent groups of Wistar rats distributed according to treatment: cocaine plus alcohol (C+A), cocaine (C), or physiological saline (S) as the control condition. The primary experimental groups were the cocaine (C) and the physiological saline (S) groups, as the main objective was to investigate the neurobiological effects of combined cocaine and alcohol self-administration.

**Fig. 1.**
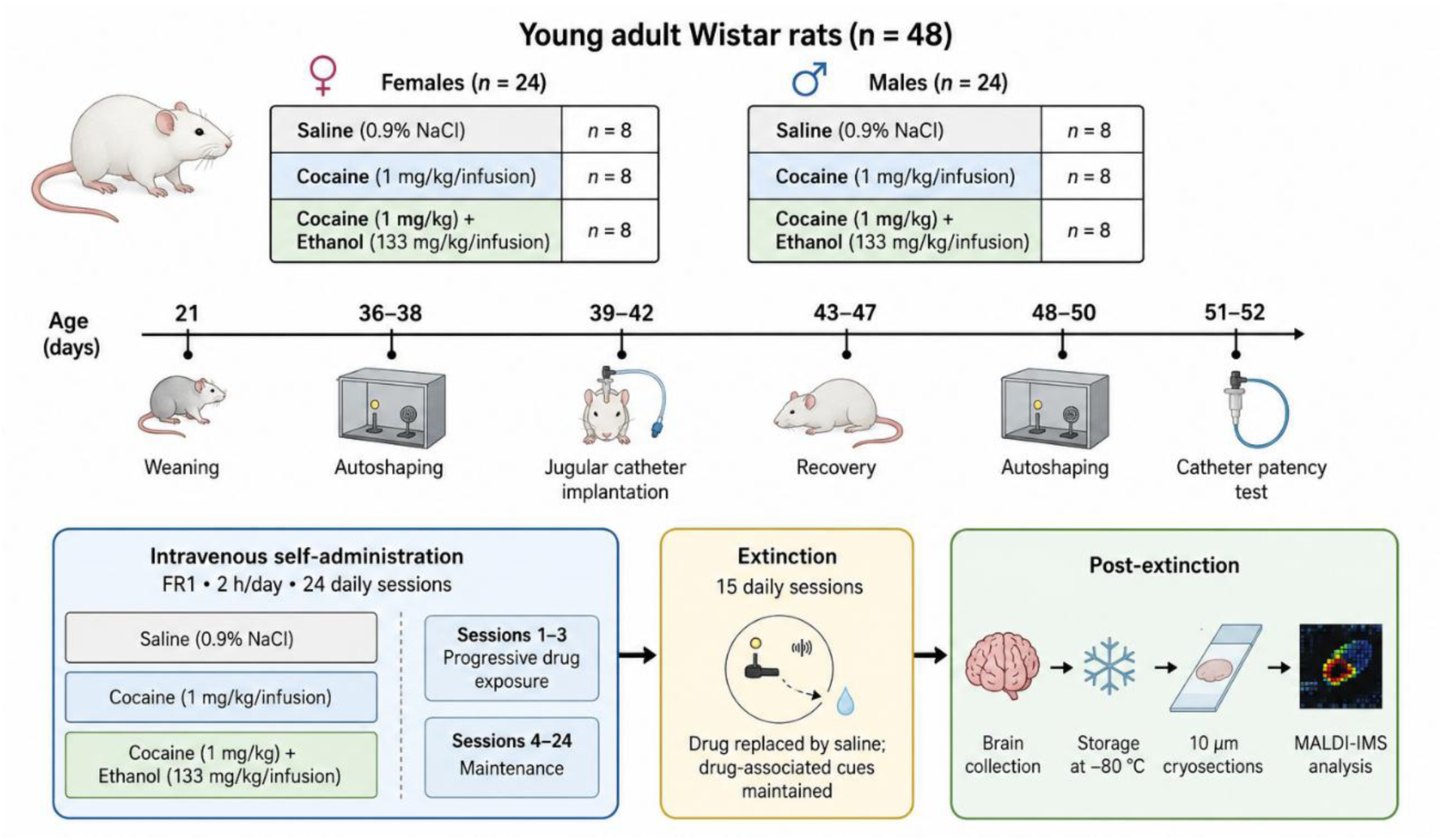
Experimental design of cocaine and combined cocaine–alcohol self-administration followed by extinction.

The cocaine (C) group was included to determine whether the molecular alterations associated with combined cocaine and alcohol self-administration were also present following cocaine self-administration alone. Alcohol alone was not included because rats do not reliably maintain intravenous alcohol self-administration under the experimental conditions employed. The number of animals was selected to ensure sufficient statistical power while adhering to the 3Rs principle.

### Surgical procedures

For intravenous drug self-administration, a polyvinyl chloride catheter (0.064″ internal diameter) was surgically implanted into the right jugular vein, with the catheter tip positioned near the right atrium. All surgical procedures were performed under isoflurane anaesthesia (5% for induction and 2% for maintenance; La Bouvet, France). Marbofloxacin (0.25 mg/kg, i.v.) was administered on the day of surgery and the following day as prophylactic antibiotic treatment. Postoperative analgesia consisted of buprenorphine (0.05 mg/kg, s.c.) administered for four consecutive days after surgery. Catheters were flushed daily with 0.5 mL of a heparin–gentamicin solution to maintain catheter patency and prevent infection. Catheter functionality was assessed 24 h before the first self-administration session and again 24 h after the last session by intravenous administration of sodium thiopental (0.10 mg/kg). Catheters were considered patent when thiopental administration produced rapid loss of consciousness.

### Drug self-administration

Intravenous self-administration was conducted in standard operant conditioning chambers (Coulbourn Instruments, USA). Responses on the active lever triggered an intravenous infusion containing cocaine (1 mg/kg) for the cocaine group, cocaine (1 mg/kg) plus ethanol (133 mg/kg) in a single solution for the combined cocaine– alcohol group, or 0.9% NaCl for control animals. Self-administration sessions were controlled using custom software based on the MED-PC platform (Med Associates, St. Albans, VT, USA). Self-administration was conducted under a fixed-ratio 1 schedule of reinforcement (FR1), under which each active lever press resulted in one infusion. Using this procedure, rats have previously been shown to acquire stable patterns of cocaine and combined cocaine–alcohol self-administration (Marcos et al., 2020). The self-administration phase lasted 24 days and consisted of daily 2-h sessions. During the first three sessions, exposure to cocaine, alone or in combination with alcohol, was gradually increased, after which animals continued under maintenance conditions until completion of the self-administration period. Subsequently, rats underwent 15 daily extinction sessions. During extinction, the drug solutions were replaced with saline; therefore, responses on the previously active lever resulted in saline infusion, while animals remained exposed to the same contextual and conditioned cues present during the acquisition and maintenance phases. Detailed behavioural procedures and self-administration data have been reported previously (Marcos et al., 2023, 2020).

The animals were sacrificed by decapitation after the final session. The brains were removed and preserved at -80 °C until further analysis.

### Dissection and tissue preparation for MALDI-IMS imaging of tryptic peptides

Brains were sectioned sagittaly at a thickness of 10 μm using a Leica CM3050 S cryostat and thaw-mounted onto conductive indium tin oxide (ITO)-coated glass slides (Bruker Daltonics). Two consecutive tissue sections were obtained from each animal and selected to include the brain regions of interest, namely the nucleus accumbens (ACC), and ventral hippocampus (vHPC), according to the rat brain atlas (Paxinos and Watson, 2017). All sections were collected from the right hemisphere. The anatomical localization and boundaries of the regions of interest are shown in **Fig. 2**, based on corresponding atlas sections.

**Fig. 2.**
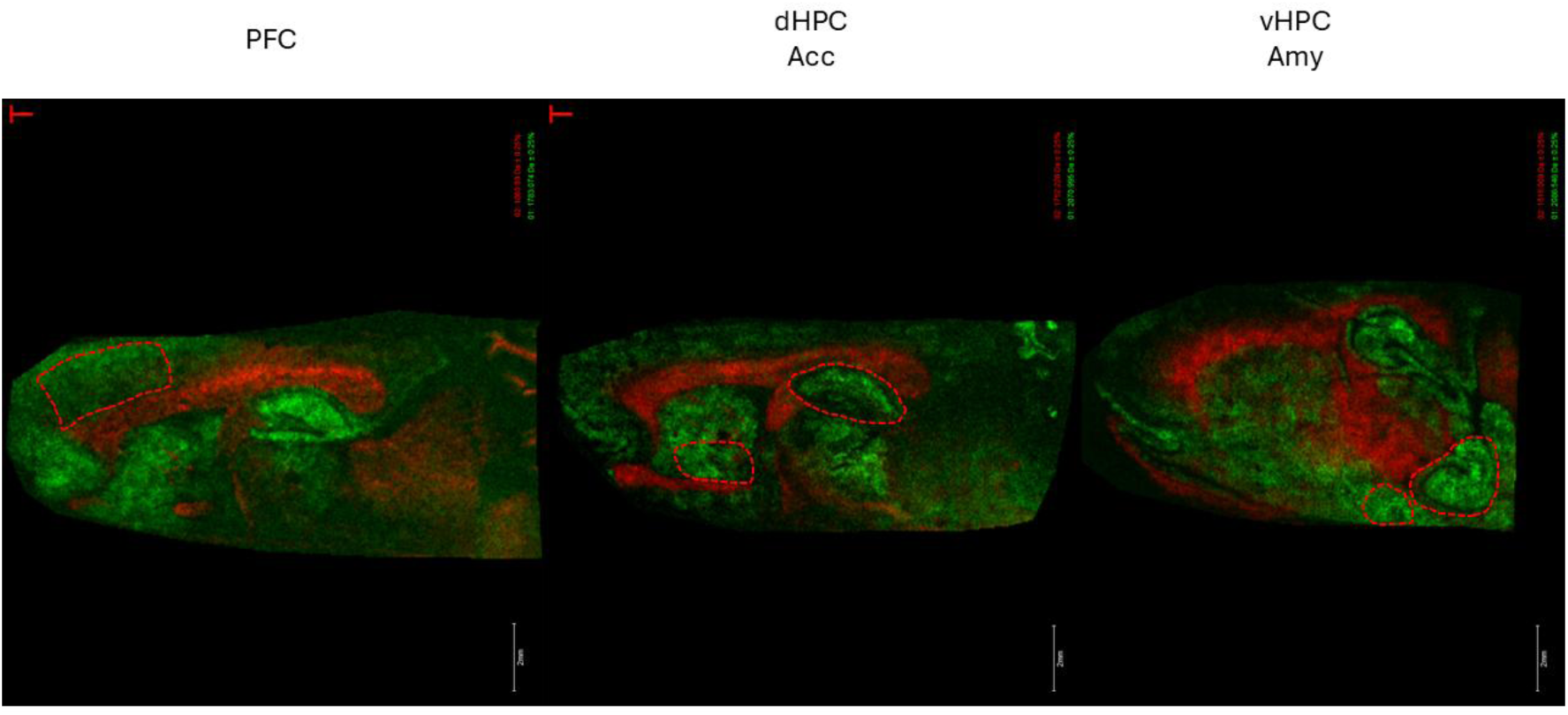
Anatomical localization of the brain regions of interest in representative sagittal rat brain sections. Red dashed lines delineate the five regions routinely evaluated in our MALDI imaging workflow: prefrontal cortex (PFC), dorsal hippocampus (dHPC), nucleus accumbens (ACC), ventral hippocampus (vHPC), and amygdala (AMY). In the present study, only the ACC and vHPC were included in the statistical and peptidomic analyses. Anatomical boundaries were established according to the rat brain atlas of Paxinos and Watson (2017). All tissue sections were obtained from the right hemisphere. Scale bar: 2 mm.

The slides were stored at room temperature until the day of the analysis. To remove most of the lipids from the sections, slices were washed in Carnoy’s solution (6 ethanol: 3 chloroform: 1 acetic acid; all from Thermo Fisher, USA) (Deutskens et al., 2011), and allowed to dry. Then, a solution containing 25 µg/mL of sequencing grade trypsin (Promega, USA) diluted in 20 mM ammonium bicarbonate (Thermo Fisher, USA) was uniformly applied onto the tissue sections using a HTXTM SprayerTM (HTX Technologies, USA) in 15 layers under the following conditions: nozzle height 40 mm, nitrogen pressure 10 psi, spray flow rate of 10 µL/min, temperature 30 °C, z-arm velocity 750 mm/min, track spacing 2 mm, crisscross pattern and drying time 0 s. Samples were then incubated overnight with trypsin in a humidity chamber at 37°C. After air-drying, the slides were coated with 7 mg/mL of α-Cyano-4-hydroxycinnamic acid (CHCA, Bruker Daltonics, Germany) in 70% acetonitrile (ACN)/0.1% trifluoroacetic acid (TFA) using the same sprayer with the following parameters: 8 passes, nozzle height 40 mm, nitrogen pressure 10 psi, flow rate of 0.1 mL/min, temperature 75°C, z-arm velocity 1200 mm/min, track spacing 3 mm, crisscross pattern and drying time 0 s. Following air-drying, the samples were immediately analysed using MALDI imaging.

### MALDI imaging acquisition and analysis

All imaging analysis were performed using a RapifleXTM MALDI TissuetyperTM TOF/TOF mass spectrometer (Bruker Daltonics, Germany) equipped with a Smartbeam™ 3D laser.

Mass measurements were performed in reflector positive ion mode in the m/z range of 600 to 3500 Da, with a spatial resolution of 50 μm. External calibration was performed using the Peptide Calibration Standard Kit II (Bruker Daltonics, Germany).

Visualization and statistical analysis of the peak list were performed with FlexImaging and SCiLS Lab software (version 2026b; Bruker Daltonics, Germany). Data from the tissue sections were imported to SCiLS Lab. The processing steps included baseline subtraction (Top-hat filter), normalization (Total Ion Current algorithm), and spatial denoising (weak). Peaks were aligned to the mean spectrum by centroid matching. Average spectra, representative of the whole measurement regions and ROIs, were generated to display differences in the peptide profiles. Intensity values for each m/z peak were exported to Excel for further calculations.

### MS/MS of digested tissue sections

Once each m/z was analysed, the precise identification of the corresponding peptide of the ones showing significant differences between the groups was performed by spatially targeting and sequencing peptides using traditional tandem mass spectrometry (MS/MS) approaches directly on the tissue. The resulting MS/MS spectra were submitted to a MASCOT (Matrix Science, USA) database search engine using BioTools software (Bruker Daltonics, Germany) to match tryptic peptide sequences to their respective intact proteins. The search was performed with a parent ion tolerance of 100-200 ppm and a fragment ion tolerance of ± 0.5 Da against the *Rattus norvegicus* database. The search criteria also included up to two missed cleavages and variable modifications, including protein N terminus acetylation, histidine/tryptophan oxidation, and methionine oxidation.

### Statistical analysis

Data were analysed by three way ANOVA, using treatment, sex and ROI as factors, and followed by Tukey’s *post-hoc* analysis. Given the high dimensionality of MALDI imaging datasets and the exploratory nature of the present study, statistical significance was established using a false discovery rate (FDR) threshold of q < 0.20 following Benjamini–Hochberg correction. This threshold is consistent with recommendations for exploratory MALDI imaging studies, where a less stringent FDR may improve sensitivity for biologically relevant feature discovery while maintaining appropriate control of multiple testing (Ngai et al., 2025; Palmer et al., 2017). Candidate m/z features were then subjected to additional selection criteria, including a minimum fold change (>1.5) and visual validation of ion images. All analyses were performed using Perseus software v.2.1.6 (Tyanova et al., 2016).

## Results

The present study investigated peptide/protein alterations induced by intravenous cocaine and cocaine-plus-alcohol self-administration using MALDI imaging mass spectrometry (MALDI-IMS). Peptide signatures were analysed in two brain regions critically involved in reward processing and addiction, the nucleus accumbens (ACC) and ventral hippocampus (vHPC), to identify treatment-dependent molecular changes associated with cocaine exposure and the modulatory effects of alcohol co-use.

### Global peptide profile differences across experimental groups

To obtain an overview of the peptide/protein profiles across experimental groups, principal component analysis (PCA) was performed independently for ACC and vHPC (**Fig. 3**). In both brain regions, the PCA revealed a tendency for samples to cluster according to treatment, although partial overlap between groups was observed.

**Fig. 3.**
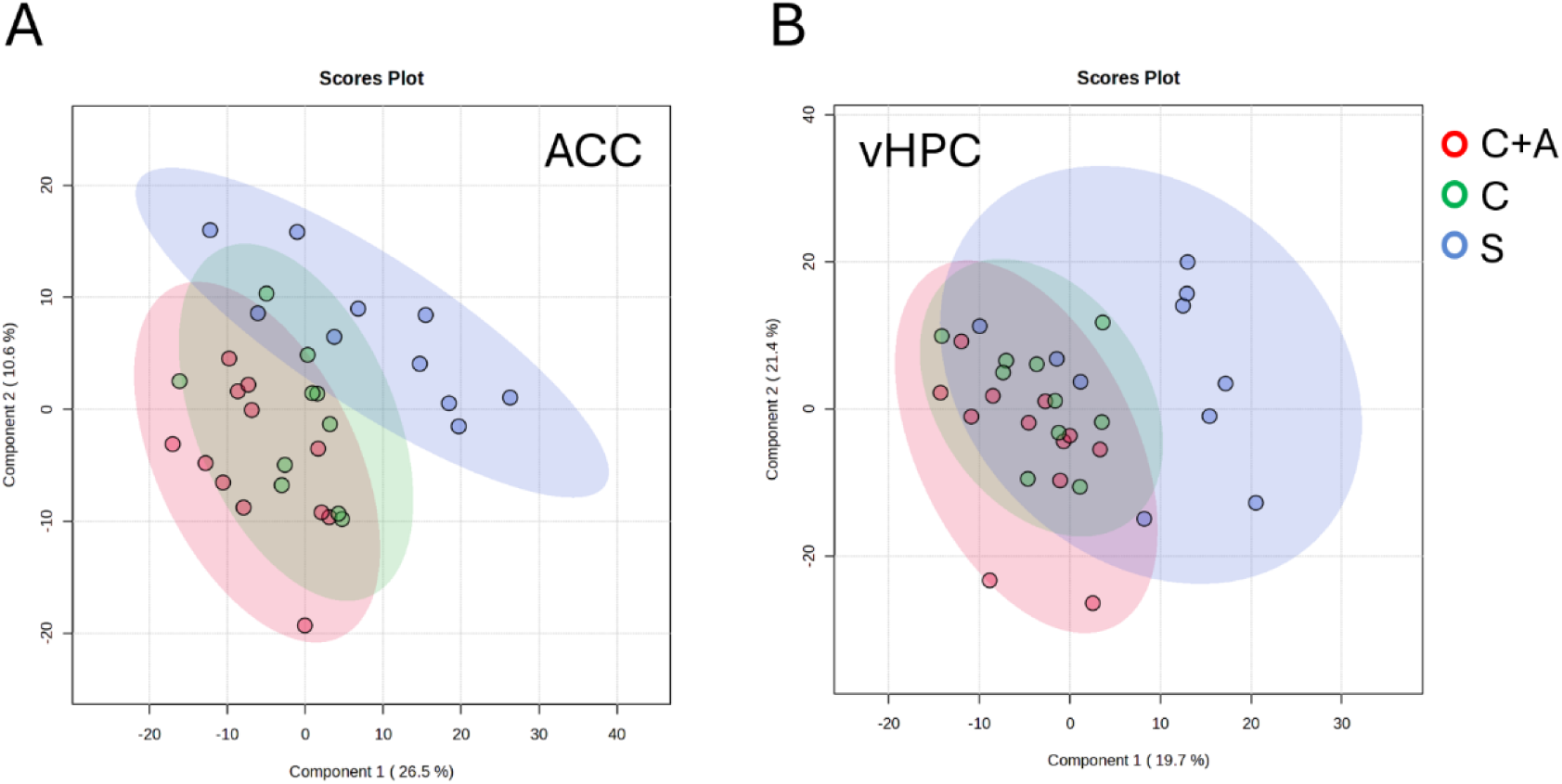
Principal component analysis (PCA) of peptide/protein profiles obtained by MALDI imaging mass spectrometry. Score plots showing the distribution of individual animals according to treatment in the (A) nucleus accumbens (ACC) and (B) ventral hippocampus (vHPC). Each point represents one animal. Shaded ellipses indicate the dispersion of each experimental group. Red, cocaine plus alcohol (C+A); green, cocaine (C); blue, saline (S).

In the ACC, the S group was clearly separated from both the C and C+A groups along the first principal component, whereas the C and C+A groups showed considerable overlap (**Fig. 3A**). A similar distribution was observed in the vHPC, where the S group formed a distinct cluster, while the C and C+A groups remained partially overlapping (**Fig. 3B**). Overall, these results suggest that drug self-administration induces broad changes in peptide/protein profiles in both brain regions, whereas alcohol co-administration modulates only part of the molecular signature associated with cocaine exposure.

### Hierarchical clustering of differentially expressed peptide/protein profiles

To further investigate treatment-dependent changes in peptide/protein expression, hierarchical clustering analysis was performed using the differentially expressed m/z features identified in the ACC and vHPC (**Fig. 4**). In both brain regions, hierarchical clustering clearly separated the S group from the drug-treated groups, whereas the C and C+A groups clustered more closely together. Similar clustering patterns were observed in both brain regions, indicating that cocaine self-administration induces marked alterations in peptide/protein expression, while alcohol co-administration preserves much of the molecular signature associated with cocaine exposure.

**Fig. 4.**
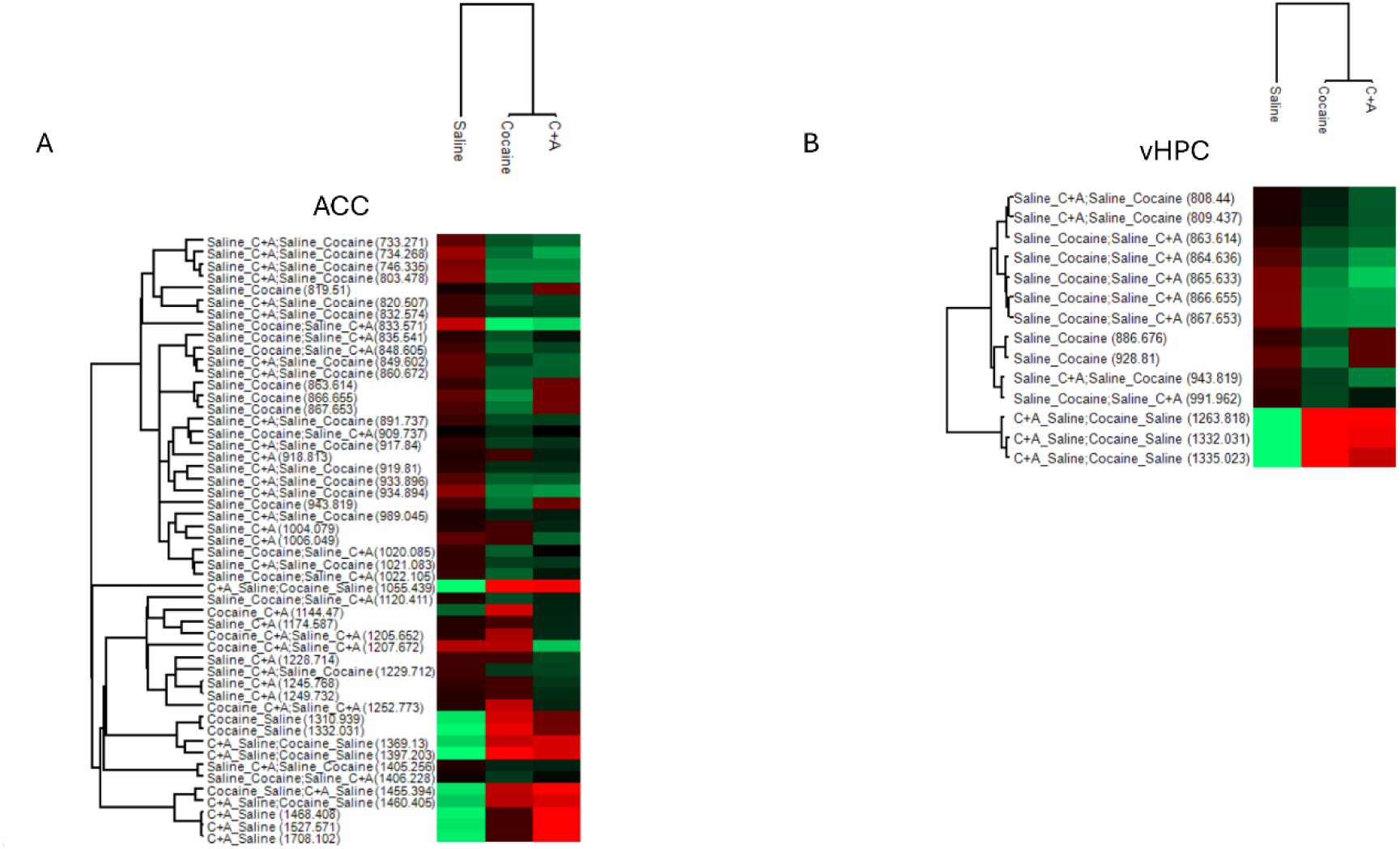
Hierarchical clustering of differentially expressed peptide/protein features identified by MALDI imaging mass spectrometry. Heatmaps showing the relative abundance of significantly altered m/z features across the experimental groups in the (A) nucleus accumbens (ACC) and (B) ventral hippocampus (vHPC). Rows represent individual peptide/protein features, identified by their m/z values, and columns correspond to the experimental groups (S, C and C+A). Hierarchical clustering was performed using Euclidean distance and complete linkage. Colour intensity represents the relative abundance of each feature, with red indicating higher relative intensity, green indicating lower relative intensity, and black representing intermediate abundance

### Differentially expressed peptide/protein features

The differentially expressed peptide/protein features identified in the ACC and vHPC are summarized in **Table 1** and **Table 2**, respectively. A total of 53 significant m/z features were detected in the ACC, whereas only 14 reached statistical significance in the vHPC, indicating a markedly greater molecular response in the ACC following cocaine and cocaine-plus-alcohol self-administration. Interestingly 5 were found in both ACC and vHPC (863.614, 866.655, 867.653, 943.819, 1332.031, **Table 1 and 2**).

**Table 1.** Differentially expressed peptide/protein features identified in the nucleus accumbens (ACC). m/z values showing significant differences among experimental groups, including −log(p) values, FDR-adjusted q-values, and significant pairwise comparisons. In the pairwise comparisons, the first group indicates the condition with the higher relative peptide intensity. Expression patterns were classified as preserved upregulation (C and C+A vs S), preserved downregulation (S vs C and C+A), reverted (significant difference between S and C but not between S and C+A), or C+A-specific (significant differences involving C+A only).

| m/z | $-\log(p)$ | q-value (FDR) | Significant comparison(s) | Expression pattern |
| --- | --- | --- | --- | --- |
| 733,271 | 2,783 | 0,074 | S vs C+A; S vs C | Preserved downregulation |
| 734,268 | 3,550 | 0,021 | S vs C+A; S vs C | Preserved downregulation |
| 746,335 | 3,345 | 0,022 | S vs C+A; S vs C | Preserved downregulation |
| 803,478 | 3,573 | 0,027 | S vs C+A; S vs C | Preserved downregulation |
| 819,510 | 1,714 | 0,192 | S vs C | Reverted |
| 820,507 | 2,478 | 0,086 | S vs C+A; S vs C | Preserved downregulation |
| 832,574 | 2,206 | 0,120 | S vs C+A; S vs C | Preserved downregulation |
| 833,571 | 4,673 | 0,012 | S vs C+A; S vs C | Preserved downregulation |
| 835,541 | 2,097 | 0,122 | S vs C+A; S vs C | Preserved downregulation |
| 848,605 | 2,610 | 0,081 | S vs C+A; S vs C | Preserved downregulation |
| 849,602 | 2,641 | 0,084 | S vs C+A; S vs C | Preserved downregulation |
| 860,672 | 2,808 | 0,079 | S vs C+A; S vs C | Preserved downregulation |
| 863,614 | 2,031 | 0,130 | S vs C | Reverted |
| 866,655 | 2,564 | 0,081 | S vs C | Reverted |
| 867,653 | 2,224 | 0,124 | S vs C | Reverted |
| 891,737 | 2,348 | 0,103 | S vs C+A; S vs C | Preserved downregulation |
| 909,737 | 1,689 | 0,196 | S vs C+A; S vs C | Preserved downregulation |
| 917,84 | 2,180 | 0,122 | S vs C+A; S vs C | Preserved downregulation |
| 918,813 | 1,798 | 0,177 | S vs C+A | C+A-specific |
| 919,81 | 2,022 | 0,129 | S vs C+A; S vs C | Preserved downregulation |

| m/z | -log(p) | q-value (FDR) | Significant comparison(s) | Expression pattern |
| --- | --- | --- | --- | --- |
| 933,896 | 2,710 | 0,080 | S vs C+A; S vs C | Preserved downregulation |
| 934,894 | 3,450 | 0,022 | S vs C+A; S vs C | Preserved downregulation |
| 943,819 | 2,154 | 0,123 | S vs C | Reverted |
| 989,045 | 1,839 | 0,170 | S vs C+A; S vs C | Preserved downregulation |
| 1004,08 | 1,767 | 0,182 | S vs C+A | C+A-specific |
| 1006,05 | 2,460 | 0,085 | S vs C+A | C+A-specific |
| 1020,09 | 2,097 | 0,118 | S vs C+A; S vs C | Preserved downregulation |
| 1021,08 | 2,211 | 0,125 | S vs C+A; S vs C | Preserved downregulation |
| 1022,11 | 2,249 | 0,123 | S vs C+A; S vs C | Preserved downregulation |
| 1055,44 | 2,524 | 0,082 | C+A vs S; C vs S | Preserved upregulation |
| 1120,41 | 2,128 | 0,122 | S vs C+A; S vs C | Preserved downregulation |
| 1144,47 | 1,747 | 0,184 | C vs C+A | Reverted |
| 1174,59 | 1,837 | 0,163 | S vs C+A | C+A-specific |
| 1205,65 | 2,004 | 0,125 | S vs C+A; S vs C | Preserved downregulation |
| 1207,67 | 3,696 | 0,034 | S vs C+A; S vs C | Preserved downregulation |
| 1228,71 | 2,131 | 0,125 | S vs C+A | C+A-specific |
| 1229,71 | 2,292 | 0,116 | S vs C+A; S vs C | Preserved downregulation |
| 1245,77 | 2,014 | 0,124 | S vs C+A | C+A-specific |
| 1249,73 | 1,871 | 0,160 | S vs C+A | C+A-specific |
| 1252,77 | 2,126 | 0,118 | C vs C+A; S vs C+A | C+A-specific |
| 1310,94 | 1,727 | 0,189 | C vs S | Reverted |
| 1332,03 | 2,034 | 0,133 | C vs S | Reverted |
| 1369,13 | 1,783 | 0,179 | C+A vs S; C vs S | Preserved upregulation |
| 1397,2 | 2,387 | 0,100 | C+A vs S; C vs S | Preserved upregulation |
| 1405,26 | 1,892 | 0,156 | S vs C+A; S vs C | Preserved downregulation |
| 1406,23 | 1,837 | 0,167 | S vs C+A; S vs C | Preserved downregulation |
| 1455,39 | 2,021 | 0,126 | C+A vs S; C vs S | Preserved upregulation |
| 1460,41 | 1,690 | 0,199 | C+A vs S; C vs S | Preserved upregulation |
| 1468,41 | 1,939 | 0,141 | C+A vs S | C+A-specific |
| 1527,57 | 1,757 | 0,184 | C+A vs S | C+A-specific |
| 1708,1 | 1,946 | 0,143 | C+A vs S | C+A-specific |

**Table 2.** Differentially expressed peptide/protein features identified in the ventral hippocampus (vHPC). m/z values showing significant differences among experimental groups, including −log(p) values, FDR-adjusted q-values, and significant pairwise comparisons. In the pairwise comparisons, the first group indicates the condition with the higher relative peptide intensity. Expression patterns were classified as preserved upregulation (C and C+A vs S), preserved downregulation (S vs C and C+A), reverted (significant difference between S and C but not between S and C+A), or C+A-specific (significant differences involving C+A only).

| m/z | $-\log(p)$ | q-value (FDR) | Significant comparison(s) | Expression pattern |
| --- | --- | --- | --- | --- |
| 808,440 | 2,385 | 0,202 | S vs C+A; S vs C | Preserved downregulation |
| 809,437 | 2,423 | 0,219 | S vs C+A; S vs C | Preserved downregulation |
| 863,614 | 2,770 | 0,120 | S vs C+A; S vs C | Preserved downregulation |
| 864,636 | 3,418 | 0,053 | S vs C+A; S vs C | Preserved downregulation |
| 865,633 | 3,975 | 0,048 | S vs C+A; S vs C | Preserved downregulation |
| 866,655 | 3,814 | 0,025 | S vs C+A; S vs C | Preserved downregulation |
| 867,653 | 3,831 | 0,036 | S vs C+A; S vs C | Preserved downregulation |
| 886,676 | 2,398 | 0,212 | S vs C | Reverted |
| 928,810 | 2,958 | 0,112 | S vs C | Reverted |
| 943,819 | 2,915 | 0,107 | S vs C+A; S vs C | Preserved downregulation |
| 991,962 | 2,371 | 0,193 | S vs C+A; S vs C | Preserved downregulation |
| 1263,818 | 3,326 | 0,056 | C+A vs S; C vs S | Preserved upregulation |
| 1332,031 | 2,812 | 0,123 | C+A vs S; C vs S | Preserved upregulation |
| 1335,023 | 2,640 | 0,141 | C+A vs S; C vs S | Preserved upregulation |

In both brain regions, most significant pairwise comparisons were observed between the S group and the drug-treated groups (C and C+A), confirming that cocaine self-administration produced widespread alterations in peptide/protein expression profiles. By contrast, only a limited number of m/z features differed between the C and C+A groups, suggesting that alcohol co-administration modified a subset of the molecular changes induced by cocaine alone.

To illustrate the main patterns of peptide/protein regulation identified in the ACC, representative violin plots of selected m/z features are shown in **Fig. 5**. Several peptide features exhibited persistent alterations following cocaine self-administration that were maintained after alcohol co-administration. For example, the m/z 833.571 showed a sustained reduction in normalized intensity in both the C and C+A groups relative to S, whereas the m/z 1397.203 displayed a persistent increase under both drug treatments (**Fig. 5A**, **Fig. 5B**). Together, these findings suggest that these molecular alterations represent core cocaine-associated responses that are relatively resistant to alcohol-induced modulation.

**Fig. 5.**
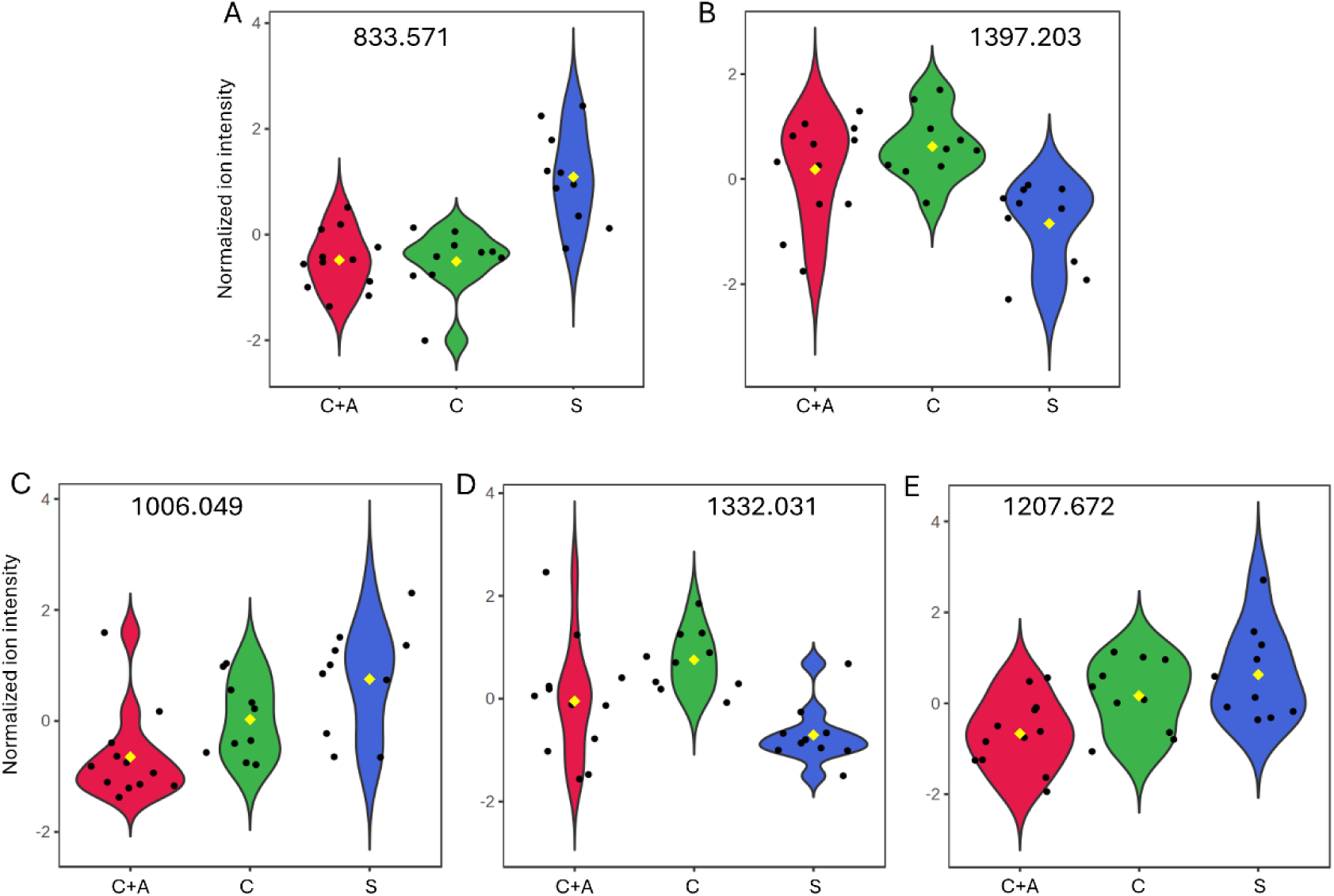
Representative violin plots showing the normalized intensity of selected m/z features identified in the nucleus accumbens (ACC). The plots illustrate representative patterns of treatment-dependent peptide/protein regulation across the experimental groups (S, C and C+A). (A) Cocaine-induced downregulated molecular feature preserved after cocaine–alcohol co-exposure. (B) Cocaine-induced upregulated molecular feature preserved after cocaine–alcohol co-exposure. (C) Molecular feature specifically altered following cocaine–alcohol co-exposure. (D) Cocaine-induced upregulated molecular feature attenuated following cocaine–alcohol co-exposure. (E) Molecular feature selectively decreased following cocaine–alcohol co-exposure.

Other peptide features displayed treatment-specific response patterns. The m/z 1006.049 was selectively altered in the C+A group, suggesting the emergence of molecular changes specifically associated with cocaine-alcohol co-use (**Fig. 5C**). Importantly, these alterations were not detected in the cocaine-only condition, suggesting that they may represent molecular adaptations specifically associated with polysubstance exposure rather than simple cocaine-induced effects. Conversely, the m/z 1332.031 exhibited an increase following cocaine self-administration that was no longer significant after alcohol co-administration, indicating an attenuation of the cocaine-induced effect (**Fig. 5D**). This may indicate that alcohol may partially normalize, counteract or mask specific molecular responses induced by cocaine exposure alone. Finally, the m/z 1207.672 showed a selective increase in the C+A group compared with both S and C, representing a peptide feature specifically associated with combined cocaine and alcohol exposure (**Fig. 5E**). These findings suggest that alcohol co-exposure does not simply introduce additional molecular changes but may actively modulate the magnitude and direction of specific cocaine-induced molecular responses within the analysed brain regions.

Representative violin plots illustrating the principal patterns of peptide/protein regulation identified in the vHPC are shown in **Fig. 6**. Similar to the ACC, several peptide features exhibited persistent cocaine-induced alterations that remained unchanged after alcohol co-administration. Thus, the m/z 867.653 showed a sustained reduction, whereas the m/z 1263.818 displayed a persistent increase in both the C and C+A groups relative to S (**Fig. 6A**, **Fig. 6B**). In contrast, the m/z 928.810 exhibited a cocaine-induced decrease that was attenuated following alcohol co-administration, suggesting that alcohol selectively modulated a subset of the molecular changes induced by cocaine (**Fig. 6C**).

**Fig. 6.**
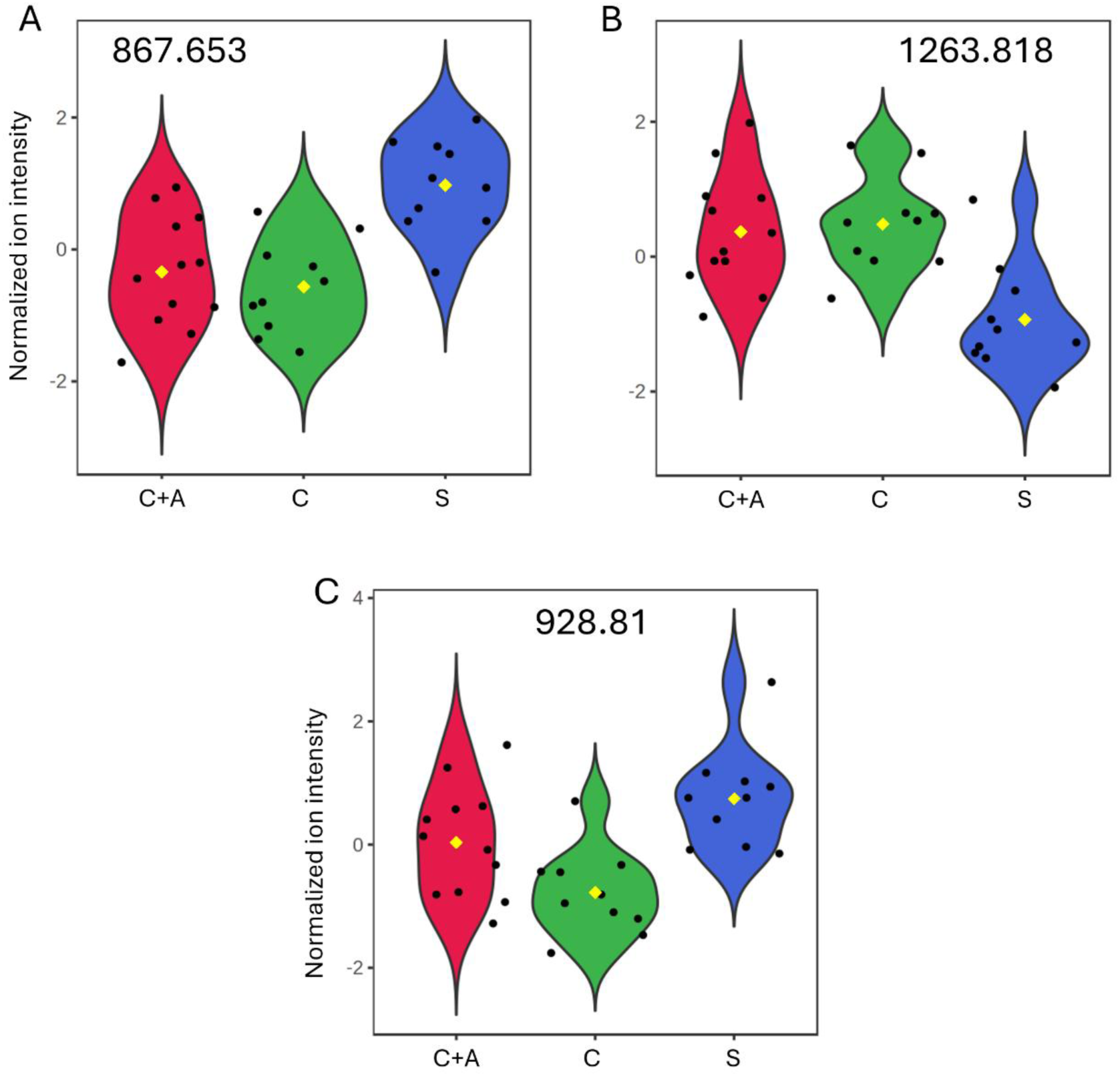
Representative violin plots showing the normalized intensity of selected m/z features identified in the ventral hippocampus (vHPC). The plots illustrate representative patterns of treatment-dependent peptide/protein regulation across the experimental groups (S, C and C+A). (A) Cocaine-induced downregulated molecular feature preserved after cocaine–alcohol co-exposure. (B) Cocaine-induced upregulated molecular feature preserved after cocaine–alcohol co-exposure. (C) Cocaine-induced downregulated molecular feature attenuated following cocaine–alcohol co-exposure.

### Putative protein identification of differential m/z features

To gain insight into the molecular identity of the differential m/z features detected by MALDI-IMS, selected ions were subjected to tandem mass spectrometry (MS/MS) analysis. Fragmentation spectra were analysed using BioTools software and searched against protein databases using Mascot to obtain putative peptide and protein identifications. Representative MS/MS spectra are shown in **Supplementary Figure S1**. Putative peptide and protein assignments for 19 selected m/z features are presented in **Table 3**, together with their BioTools scores and fragmentation p-values. All reported assignments showed significant fragmentation matches (p < 0.05).

**Table 3.** Proteins identified by MALDI-MS/MS from significant m/z features detected in the nucleus accumbens (ACC) and ventral hippocampus (vHPC). Protein identification was performed using BioTools software and Mascot database searching. Representative function summarizes the principal biological role of each protein. Expression pattern describes the regulation profile observed in each brain region and was classified as preserved upregulation, preserved downregulation, reverted, or C+A-specific according to the significant pairwise comparisons among the S, C, and C+A groups. The BioTools score reflects the quality of the match between the experimental MS/MS spectrum and the assigned peptide, whereas the fragmentation p-value indicates the statistical significance of the peptide identification.

| m/z | Brain region | Protein | Full name | Representative biological process | Expression pattern | Biotoools score | Fragmentation p-value |
| --- | --- | --- | --- | --- | --- | --- | --- |
| 866,655 | ACC, vHPC | **SPTN1** | Spectrin alpha chain, non-erythrocytic 1 | Membrane stability, including in neurons | Reverted (ACC); Preserved downregulation (vHPC) | 6168 | 0,006 |
| 943,819 | ACC, vHPC | **GRIN3B** | Glutamate ionotropic receptor NMDA type subunit 3B | Glutamatergic signaling in neurons | Reverted (ACC); Preserved downregulation (vHPC) | 447 | 0,046 |
| 1004,079 | ACC | **CABIN1** | Calcineurin-binding protein cabin-1 | Calcium regulation | C+A-specific | 1575 | 0,0068 |
| 1055,439 | ACC | **PLEKHM1** | Pleckstrin homology domain-containing family M member 1 | Autophagosome fusion | Preserved upregulation | 4566 | 0,0058 |
| 1174,587 | ACC | **TMEM199** | Transmembrane protein 199 | V-ATPase involvement in neurotransmission | C+A-specific | 8470 | 0,00079 |
| 1205,652 | ACC | **RTN3** | Reticulon-3 | Neurite organization | Preserved downregulation | 895 | 0,02 |
| 1207,672 | ACC | **ECE1** | Endothelin-converting enzyme 1 | Peptide processing and endothelin production | Preserved downregulation | 8192 | 0,0012 |
| 1249,732 | ACC | **ZFR** | Zinc finger RNA-binding protein | Expression Regulator | C+A-specific | 4399 | 0,00058 |
| 1252,773 | ACC | **MACROH2A1** | Core histone macro-H2A.1 | Epigenetic regulation | C+A-specific | 6284 | 0,0038 |
| 1310,939 | ACC | **RERE** | Arginine-glutamic acid dipeptide repeats protein | Nervous system development | Reverted | 8294 | 3,60E-06 |
| 1332,031 | ACC, vHPC | **KIF22** | Kinesin family member 22 | Microtubule organization | Reverted (ACC); Preserved upregulation (vHPC) | 898 | 0,00021 |
| 1369,13 | ACC | **MYL1** | Myosin light chain 1 | Myosin-related processes | Preserved upregulation | 12305 | 0,042 |
| 1397,203 | ACC | **HRH3** | Histamine receptor H3 | Histaminergic neurons in the hypothalamus, projected to other brain regions | Preserved upregulation | 6182 | 0,0082 |
| 1406,228 | ACC | **SLC30A3** | Zinc transporter 3 | Zinc transporter detected in neurons | Preserved downregulation | 33359 | 0,067 |
| 1455,394 | ACC | **DAGLB** | Diacylglycerol lipase beta | Endocannabinoid system | Preserved upregulation | 4364 | 0,025 |
| 1460,405 | ACC | **HMGN2** | High mobility group nucleosomal-binding domain-containing protein 2 | Chromatin remodeling | Preserved upregulation | 3079 | 0,039 |
| 1468,408 | ACC | **SLC44A3** | Choline transporter-like protein 3 | Choline transport and membrane metabolism | C+A-specific | 897 | 0,0039 |
| 1527,571 | ACC | **NDUFV3** | NADH dehydrogenase [ubiquinone] flavoprotein 3 | Electron transport chain | C+A-specific | 8260 | 0,001 |
| 1708,102 | ACC | **NOSTRIN** | Nitric oxide synthase trafficking inducer | Regulation of nitric oxide signaling | C+A-specific | 17151 | 0,00036 |

The putative protein identifications were associated with several biological processes relevant to neuronal function. These included proteins involved in neurotransmission and synaptic signalling (GRIN3B, HRH3, SLC30A3, DAGLB and ECE1), cytoskeletal organization and intracellular transport (SPTN1, RTN3, KIF22 and MYL1), chromatin remodelling and gene regulation (MACROH2A1, HMGN2, ZFR and RERE), as well as proteins related to calcium homeostasis (CABIN1), autophagy (PLEKHM1), membrane trafficking (TMEM199), choline metabolism (SLC44A3), mitochondrial respiration (NDUFV3) and nitric oxide signalling (NOSTRIN).

Among the identified proteins, three were detected in both the ACC and vHPC: spectrin alpha chain, non-erythrocytic 1 (SPTN1), glutamate ionotropic receptor NMDA type subunit 3B (GRIN3B), and kinesin family member 22 (KIF22). These shared identifications suggest that some molecular alterations induced by cocaine and cocaine–alcohol exposure are conserved across both brain regions, whereas the remaining identified proteins were region-specific and were exclusively detected in the ACC.

## Discussion

This exploratory study characterized spatial peptide/protein alterations persisting after extinction of drug-seeking behaviour in rats previously exposed to cocaine or combined cocaine–alcohol self-administration. MALDI imaging enabled the identification of treatment-related molecular patterns across brain regions involved in addiction. The objective of the present work was not to validate predefined molecular targets but to identify candidate peptide/protein alterations associated with cocaine–alcohol polysubstance use. Accordingly, an FDR threshold of 0.20 was adopted, in line with previous discovery-based omics studies, to maximize sensitivity while maintaining control of multiple testing. Importantly, all selected molecular features underwent subsequent MS/MS analysis, providing an additional level of validation beyond the initial statistical screening.

The study focused on the anterior cingulate cortex (ACC) and ventral hippocampus (vHPC) due to their well-established involvement in addiction-related corticolimbic circuitry. The ACC was prioritized because it exhibited the most prominent and consistent molecular alterations across experimental conditions, supporting its central role in executive control, craving and compulsive drug-seeking behaviours. The vHPC was included as a complementary region because of its contribution to contextual memory, emotional processing and relapse-associated mechanisms linked to substance abuse.

The substantially higher number of altered molecular features detected in the ACC suggests that this region may be particularly sensitive to the persistent neurobiological effects of cocaine and combined cocaine–alcohol exposure. The ACC is a key component of addiction-related circuitry and contributes to salience processing, motivation, inhibitory control and decision-making. Through its extensive connections with corticolimbic regions, including the nucleus accumbens, amygdala, hippocampus and medial prefrontal cortex, the ACC integrates reward-related, emotional and executive processes that may contribute to the broader molecular alterations observed in this region compared with the vHPC (Zhao et al., 2020).

The molecular alterations shared by the cocaine and cocaine–alcohol groups suggest the persistence of a core set of neuroadaptations primarily driven by cocaine exposure. Cocaine self-administration is known to induce long-lasting molecular and synaptic alterations that can persist during abstinence and after extinction training (Jarvis et al., 2019; Siciliano et al., 2016). In the present study, the maintenance of these molecular features following alcohol co-exposure suggests that some cocaine-induced responses are relatively stable and only modestly influenced by alcohol.

The molecular features exclusively altered in the cocaine–alcohol group suggest that combined exposure produces a distinct molecular profile rather than merely reproducing the effects of cocaine alone. This interpretation is consistent with previous studies showing that cocaine and alcohol co-use induces specific neuroadaptations and modifies glutamatergic signalling compared with cocaine self-administration alone (Stennett and Knackstedt, 2020). Similar treatment-specific peptide/protein alterations have also been reported using MALDI imaging mass spectrometry in other brain regions following combined cocaine and alcohol administration by our collaborative research consortium (Marcos et al., 2025). These features may therefore represent candidate molecular signatures associated with cocaine–alcohol co-exposure.

Conversely, several cocaine-associated molecular alterations were no longer detected following alcohol co-exposure, while other features differed directly between the cocaine and cocaine–alcohol groups. These findings indicate that alcohol does not simply potentiate cocaine-induced effects but can selectively reshape specific molecular responses. This interpretation is consistent with previous studies showing that cocaine–alcohol exposure produces neuroadaptations distinct from those induced by cocaine alone, including differential alterations in glutamate homeostasis and cortical responses (Mesa et al., 2023; Stennett et al., 2020).

Although fewer molecular alterations were detected in the vHPC than in the ACC, this region remains highly relevant to addiction-related neuroadaptations because of its role in contextual memory, emotional regulation and relapse. The vHPC forms part of the corticolimbic circuitry underlying substance use disorders through its reciprocal connections with the nucleus accumbens, amygdala and medial prefrontal cortex, where it contributes to the integration of contextual and emotional information that drives drug-seeking behaviour. Previous experimental studies have shown that cocaine exposure induces persistent functional and synaptic alterations within the ventral hippocampus, contributing to cocaine-associated memory and reinstatement of drug seeking (Lasseter et al., 2010; Zhou et al., 2019). Accordingly, the molecular changes identified in this region likely reflect complementary neuroadaptations associated with cocaine and cocaine–alcohol exposure.

Overall, the greater number of altered molecular features detected in the ACC suggests a broader molecular response to cocaine exposure in this region, whereas the vHPC exhibited a more restricted but treatment-specific profile. This regional divergence may reflect their distinct functional roles within addiction-related circuitry, with the ACC primarily involved in salience processing and executive control, and the vHPC contributing more selectively to contextual and emotional aspects of drug-associated memory.

The present study was designed as a discovery-based analysis aimed at identifying treatment-associated molecular patterns rather than validating individual peptide or protein targets. Consequently, the biological significance of the detected molecular features should be interpreted cautiously until their identities and functional relevance are confirmed in complementary studies. Nevertheless, the consistent regional and treatment-dependent patterns observed support the utility of MALDI imaging mass spectrometry as an unbiased approach for identifying candidate molecular pathways involved in cocaine and cocaine–alcohol-induced neuroadaptations. Future studies integrating these spatial molecular profiles with complementary proteomic analyses and independent experimental cohorts will be essential to validate the identified candidates and determine their contribution to the neurobiological mechanisms underlying cocaine and cocaine–alcohol polysubstance use.

## Acknowledgements

The authors would like to thank the following laboratory technicians for their valuable help in carrying out this study: Pilar Alberdi, from the School of Medicine of the University of Castilla-La Mancha; and Rosa Ferrado from the Department of Psychobiology, School of Psychology, National University for Distance Learning.

## Declarations

### Declaration of generative AI and AI-assisted technologies in the manuscript preparation process

During the preparation of this work, the authors used ChatGPT (OpenAI) to assist with language editing, text refinement, formatting, and consistency checks. After using this tool, the authors carefully reviewed, edited, and validated all content as necessary and take full responsibility for the content of the published article.

### Ethics statement

All experiments were performed in accordance with European Union Laboratory Animal Care Standards (2010/63/EU) and with the approval of the Bioethics Committee of the UNED.

### Confficts of interest

The authors declare no conflicts of interest.

### Data Availability Statement

The data to support the findings of this study are available from the corresponding author upon reasonable request.

