## Supplementary material for "Exploratory spatial peptidomic profiling during incubation of drug seeking following cocaine plus alcohol self-administration in young adult rats": Suppl. Fig 1

### Mascot Search Results

#### Peptide View

MS/MS Fragmentation of **QVSTLINSTDK**
Found in **ECE1_RAT** in **SwissProt**, Endothelin-converting enzyme 1 OS=Rattus norvegicus OX=10116 GN=Ece1 PE=1 SV=2

Match to Query 1: 1204.643169 from(1205.650445,1+) index(0)
Data file DATA.TXT

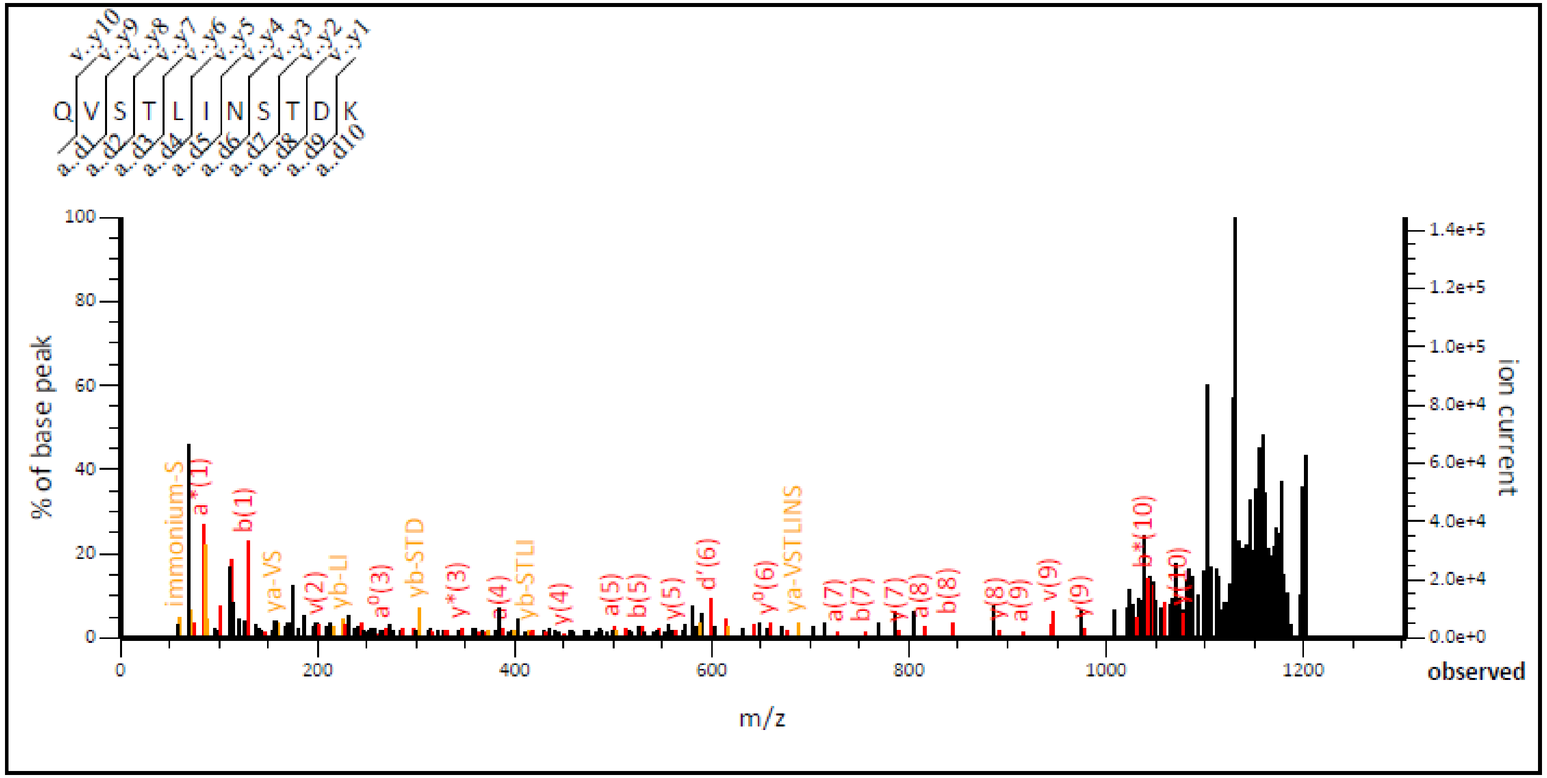


0 to 1302.72

**Monoisotopic mass of neutral peptide Mr(calc):** 1204.6299
**Ions Score:** 42 **Expect:** 0.0012
**Peak matches:** 54/194 fragment ions using 100 most intense peaks
**Annotated fragments:** 94/194 (help)

### Mascot Search Results

#### Peptide View

MS/MS Fragmentation of GNTISPIIKLLK
Found in PUR1_RAT in SwissProt, Amidophosphoribosyltransferase OS=Rattus norvegicus OX=10116 GN=Ppat PE=1 SV=1

Match to Query 1: 1295.914143 from(1296.921419,1+) index(0)
Data file DATA.TXT

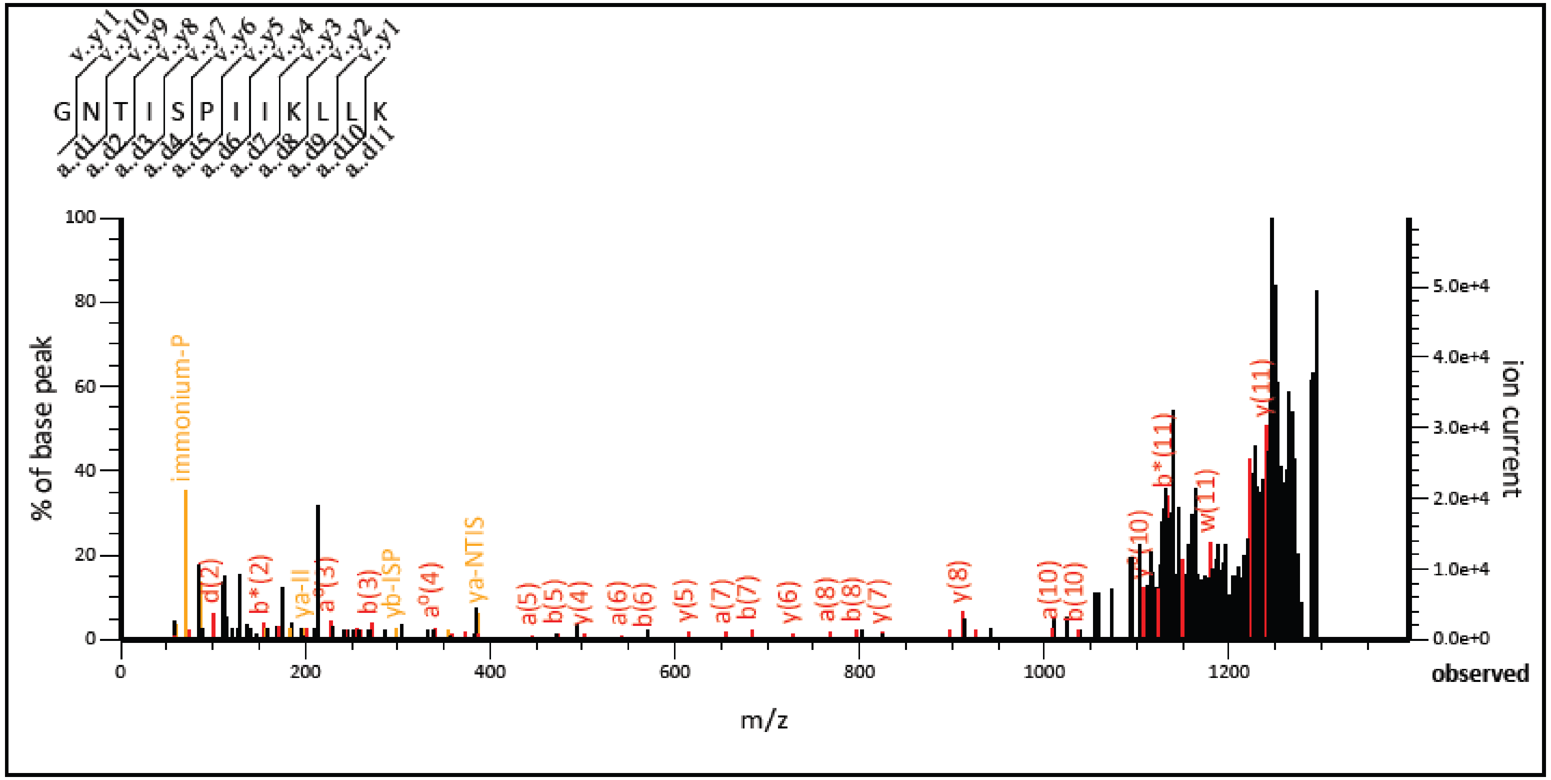


0 to 1393.94

```
Monoisotopic mass of neutral peptide Mr(calc): 1295.8176
Ions Score: 39   Expect: 0.00025
Peak matches: 39/208 fragment ions using 77 most intense peaks
Annotated fragments: 67/208    (help)
```

### Mascot Search Results

#### Peptide View

MS/MS Fragmentation of **LNQRSSRSHAVLLVK**
Found in **KIF22_RAT** in **SwissProt**, Kinesin-like protein KIF22 OS=Rattus norvegicus OX=10116 GN=Kif22 PE=2 SV=1

Match to Query 1: 1707.095331 from(1708.102607,1+) index(0)
Data file DATA.TXT

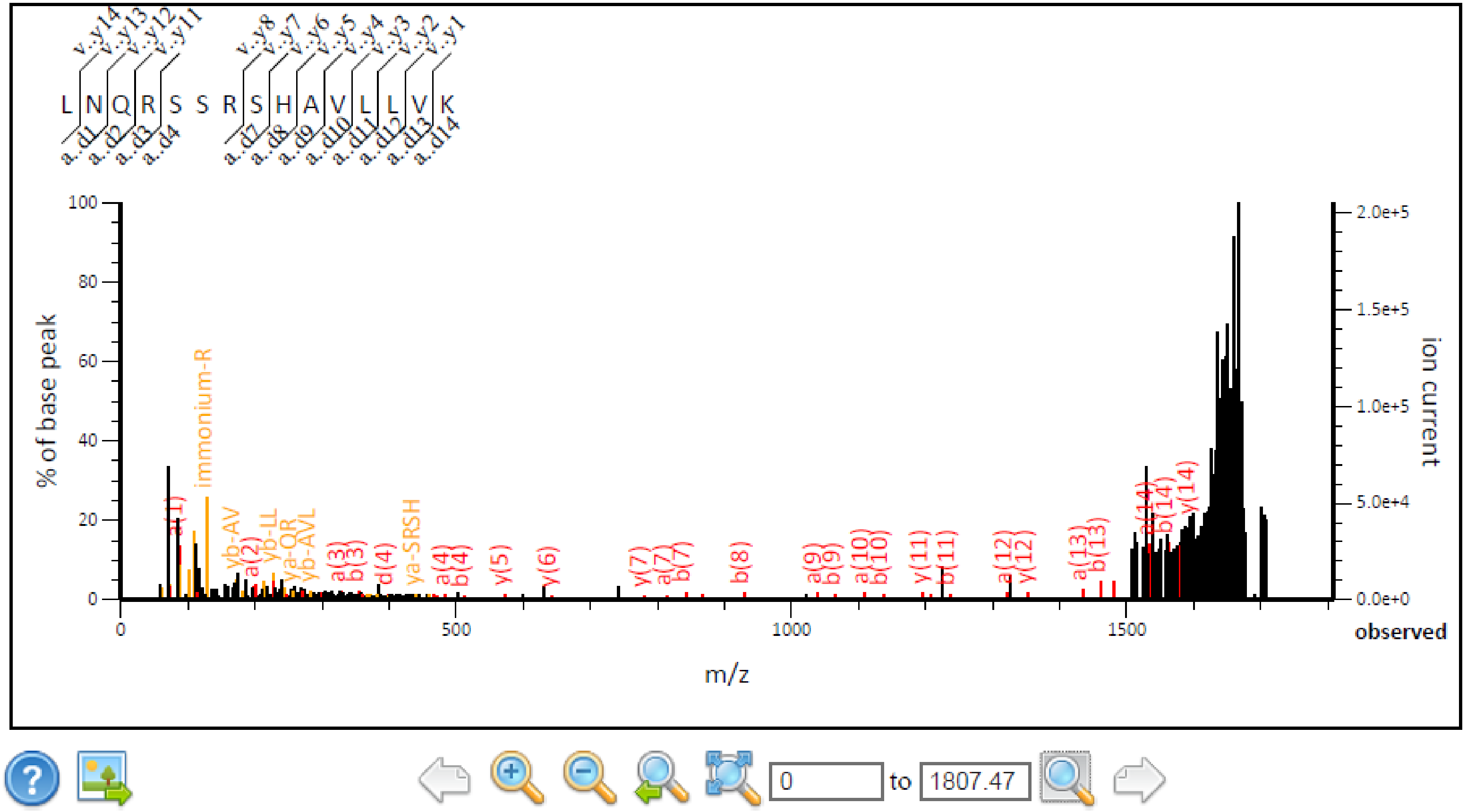


Monoisotopic mass of neutral peptide Mr(calc): 1706.9904
Ions Score: 47 Expect: 0.00021
Peak matches: 48/260 fragment ions using 78 most intense peaks
Annotated fragments: 90/260 (help)

### Mascot Search Results

#### Peptide View

MS/MS Fragmentation of **TDHFNNARSQGHK**
Found in **CTL3_RAT** in **SwissProt**, Choline transporter-like protein 3 OS=Rattus norvegicus OX=10116 GN=Slc44a3 PE=2 SV=1

Match to Query 1: 1526.550795 from(1527.558071,1+) index(0)
Data file DATA.TXT

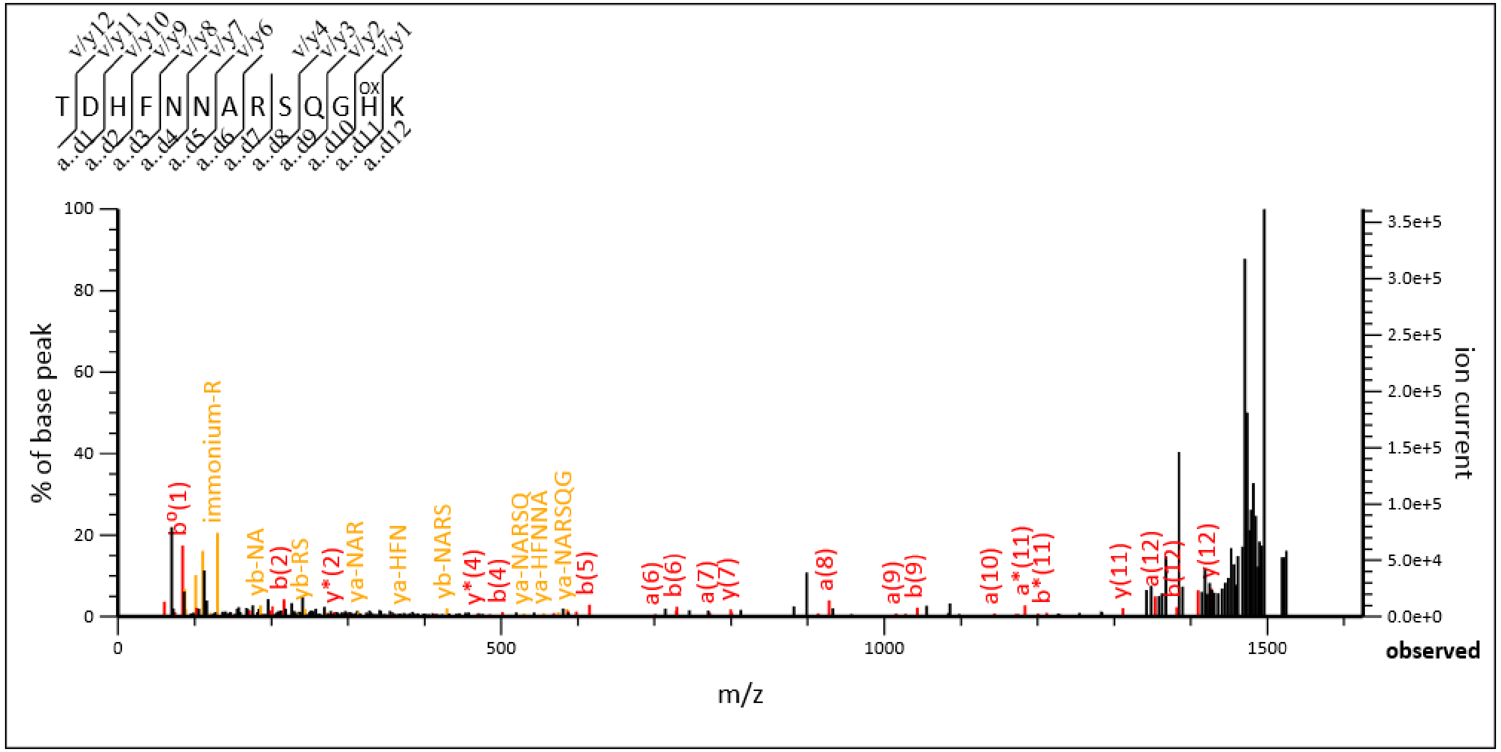


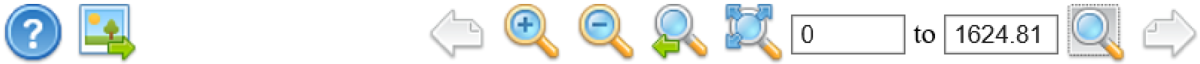

**Monoisotopic mass of neutral peptide Mr(calc):** 1526.6974
**Variable modifications:**
H12 : Oxidation (HW)
**Ions Score:** 29 **Expect:** 0.0039
**Peak matches:** 46/213 fragment ions using 110 most intense peaks
**Annotated fragments:** 75/213 (help)
